# Intestinal epithelial *Atg16l1* influences pregnancy-induced fecal microbiota shifts in mice

**DOI:** 10.1101/2023.12.10.570427

**Authors:** Víctor A. López-Agudelo, Maren Falk-Paulsen, Ateequr Rehman, Richa Bharti, Felix Sommer, Eike Matthias Wacker, David Ellinghaus, Anne Luzius, Laura Sievers, Arthur Kaser, Philip Rosenstiel

## Abstract

Throughout gestation, the female body undergoes a series of transformations, including profound alterations in intestinal microbial communities. Changes gradually increase towards the end of pregnancy and comprise reduced α-diversity of microbial communities and an increased propensity for energy harvest. Despite the importance of the intestinal microbiota for the pathophysiology of inflammatory bowel diseases, very little is known about the relationship between these microbiota shifts and pregnancy-associated complications of the disease. Here, we explored the longitudinal dynamics of gut microbiota composition and functional potential during pregnancy and after lactation in *Atg16l1^ΔIEC^*mice carrying an intestinal epithelial deletion of the Crohńs disease risk gene *Atg16l1.* Using 16S rRNA amplicon and shotgun metagenomic sequencing, we demonstrated divergent temporal shifts in microbial composition between *Atg16l1* wildtype and *Atg16l1^ΔIEC^*pregnant mice in trimester 3, which was validated in an independent experiment. Observed differences included microbial genera implicated in IBD such as *Lachnospiraceae*, *Roseburia*, *Ruminococcus*, and *Turicibacter*. Changes partially recovered after lactation. In addition, functional inference of metagenomic data suggest a reduced potential to biosynthesize mucosal protective polyamines and reduced capacity to metabolize acidic polysaccharides (ketogluconate metabolism).

On the host side, we found that the immunological response of *Atg16l1^ΔIEC^* mice is characterized by higher colonic mRNA levels of TNFA, and CXCL1 in trimester 3 and a lower weight of offspring at birth. Understanding pregnancy-dependent microbiome changes in the context of IBD may constitute the first step in the identification of fecal microbial biomarkers and microbiota-directed therapies that could help improving precision care for managing pregnancies in IBD patients.

## Introduction

Pregnancy is a transient physiological state characterized by significant alterations in the body’s physiology. These changes encompass hormonal fluctuations, substantial shifts in energy metabolism (Lain and Catalano, 2007; Mor and Cardenas, 2010; Abu-Raya *et al*., 2020) as well as changes of the immunological network state to safeguard the developing fetus thereby maximizing reproductive success. Evidence has emerged suggesting that variation in the composition and function of the intestinal microbiota is contributing to this physiological adaptation, e.g. by increasing energy harvest from nutrient intake. (Koren *et al*., 2012; Ma *et al*., 2014; Prince *et al*., 2014; DiGiulio *et al*., 2015; Edwards *et al*., 2017; Ferrocino *et al*., 2018; Faas *et al*., 2020; Kimura *et al*., 2020; Yang *et al*., 2020). Reports have highlighted bacterial community shifts between the first and third trimester, with third-trimester fecal microbiota exerting a subtle pro-inflammatory tone, which may contribute to higher adiposity and insulin insensitivity (Collado *et al*., 2008; Koren *et al*., 2012; Ferrocino *et al*., 2018). Yet, the exact mechanisms and the impact of these changes on pregnancy-induced immune alterations are still unclear (De Siena *et al*., 2021).

Despite improved therapies optimal clinical care for female patients with chronic inflammatory bowel disease during and after pregnancy remains a challenge. Data on disease course modulation by pregnancy in IBD is complex, with a significant burden resulting from pregnancy complications and relapses during or after pregnancy (Brondfield and Mahadevan, 2023). Recent evidence has shown that 30% of patients with IBD will suffer from flaring disease in the first half year after delivery despite continued immunosuppressive therapy. Given the significant role of the intestinal microbiota for most facets of IBD, it is tempting to speculate that microbial shifts during pregnancy could also contribute to the heterogeneous outcome of pregnancy and disease course modulation around pregnancy and lactation. While a larger study in 358 pregnant IBD patients from Mount Sinai suggested a correlation of specific bacterial taxa with maternal fecal calprotectin levels (Kim *et al*., 2021), no studies exist that causally link specific changes of the gut microbiome during pregnancy to inflammatory reactions or vice versa IBD-related changes of intestinal microbes to pregnancy outcomes.

In the past decade >200 IBD risk loci have been identified by genome wide association studies and exome sequencing (Jostins *et al*., 2012), that cluster in distinct molecular pathways, including autophagy (Hampe *et al*., 2007), ER stress signaling, and innate immune sensing (Franke *et al*., 2010; Jostins *et al*., 2012). Loss-of-function variants in autophagy genes, such as *ATG16L1* only affect CD patients and have been associated with clear functional defects of intestinal epithelial cells (IECs), e.g., a Paneth cell deficiency (Cadwell *et al*., 2008). Inadequate autophagy mediated by ATG16L1, via the CD T300A risk allele, increases the vulnerability of epithelial cells to inflammation induced by bacteria and viruses (Cadwell *et al*., 2010; Lassen *et al*., 2014). *Atg16l1* deficiency in the intestinal epithelium of mice leads to an age-dependent onset of subtle inflammatory changes in the ileal mucosa (Tschurtschenthaler *et al*., 2017), as well as to a paradoxical inflammatory response to the normally pro-regenerative cytokine IL-22 (Aden *et al*., 2018). Autophagy is intricately connected to the unfolded protein response (UPR) originating from the endoplasmic reticulum (Kaser *et al*., 2008). The significance of this interplay is underscored by the discovery that mice with a dual deficiency in the UPR transcription factor gene *Xbp1* and *Atg16l1* in the intestinal epithelium exhibit spontaneous transmural and fistulizing ileal inflammation resembling human Crohn’s disease (Adolph *et al*., 2013). Despite subtle findings, such as a transmissible susceptibility to DSS-induced colitis in mice with an IEC-specific deletion of the gene (Tschurtschenthaler *et al*., 2017), the role of *Atg16l1* in regulating the microbiome remains unclear.

We here hypothesized that a dysfunction of *Atg16l1* in IECs-due its altered anti-microbial effector function-could perturb the normal shift of fecal microbiota normally observed during pregnancy. We speculated that the intestinal microbiota changes during and after pregnancy could modulate susceptibility to intestinal inflammatory responses and modulate pregnancy outcomes. In this longitudinal study, we investigated the fecal microbial and physiological responses to pregnancy in mice carrying an *Atg16l1*-deletion in the intestinal epithelium and their corresponding littermate controls. We analyzed the temporal dynamics of the microbiota through 16S rRNA gene profiling and shotgun metagenomics, aiming to correlate these findings with observed microbial and immunological responses in the host. Our study provides a comprehensive view on changes in the microbial diversity during and after pregnancy. Notably, we highlight the shifting microbial composition and metabolic potential across pregnancy phases in *Atg16l1^fl/fl^* and *Atg16l1*^ΔIEC^ mice.

## Materials and Methods

### Generation and housing of conditional knockout mice

The conditional *Atg16l1* knock-out mouse line was described in detail elsewhere (Adolph *et al*., 2013; Aden *et al*., 2018). In brief, deletion of the exon 1 flanked by loxP sites was achieved by villin-promoter-driven cre-recombinase excision, which targets all major splice variants of the gene. The details and the model were described before (Adolph *et al*., 2013; Aden *et al*., 2018). Control female loxed animals (*Atg16l1^fl/fl^*) (also termed control animals or WT) were employed as littermate controls to the female villin-cre (+) mice carrying a conditional deletion in the intestinal epithelium (termed *Atg16l1^ΔIEC^* or KO). Timed matings were performed using littermate male *Atg16l1^fl/fl^*mice. The mice were housed under specific-pathogen-free (SPF) conditions in individually ventilated cages (IVCs) under a 12 h light-dark cycle. Food and water were provided ad libitum and environmental conditions were maintained at 21 °C ± 2 °C with 60 % ± 5 % humidity. All animal experiments were approved by the local animal safety review board of the federal ministry of Schleswig Holstein and conducted according to national and international laws and policies (V 241 - 69128/2016 (3-1/17) and V 242-7224.121-33). Mice were sacrificed by cervical dislocation. For RNA extraction, tissues were snap-frozen in liquid nitrogen and stored at - 80 °C. For immunohistochemistry, tissues were fixed in 10 % formalin at 4 °C for at least 24 h. For histological analysis, intestinal organs were cut longitudinally after sacrificing the mice and were stained using H&E staining, and later analyzed under the microscope. The fecal pellets sampled for reported time points were stored at -80°C until DNA extraction.

### RNA isolation and Quantitative real-time PCR analysis

For extracting tissue-specific RNA, tissue samples were mixed with 350-600 µl lysis buffer and were disrupted by rapid agitation in a TissueLyser system. Following this, the lysate was processed using the described protocol of the RNeasy Mini kit (Qiagen, Germany). Finally, the purified RNA was eluted and stored at -80°C till further usage. Expression of candidate genes (TNFA, CXCL1 and IL-10) was normalized using housekeeping genes (GAPDH/β-actin) and an unpaired non-parametric Mann-Whitney test was performed to test genotype-specific differences between nulliparous and pregnant mice as described (Aden *et al*., 2018).

### Stool DNA isolation and 16S rRNA amplicon and shotgun sequencing

DNA from stool samples from the main (87 samples) and validation (56 samples) experiments were isolated using the DNeasy PowerSoil Pro Kit (Qiagen) following the manufacturer’s protocol. Extracted DNA was eluted from the spin filter silica membrane with 100 µl of elution buffer and stored at -80 °C.

The V3-V4 region of the 16S gene was amplified using the dual barcoded primers 341F (GTGCCAGCMGCCGCGGTAA) and 806R (GGACTACHVGGGTWTCTAAT). Each primer contained additional sequences for a 12 base Golay barcode, Illumina adaptor, and a linker sequence (Caporaso *et al*., 2012). PCR was performed using the Phusion Hot Start Flex 2X Master Mix (NEB) in a GeneAmp PCR system 9700 (Applied Biosystems) and the following program [98°C for 3 min, 25x (98°C for 20 s, 55°C for 30 s, 72°C for 45 s), 72°C for 10 min, hold at 4°C]. Performance of the PCR reactions was checked using agarose gel electrophoresis. Normalization was performed using the SequalPrep Normalization Plate Kit (Thermo Fisher Scientific, Darmstadt, Germany) following the manufacturer’s instructions. Equal volumes of SequalPrep-normalized amplicons were pooled and then sequenced on an Illumina MiSeq (2 x 300 nt). Shot-gun metagenomics sequencing was carried out on DNA extracts obtained from stool samples from the main (72 samples) experiment. DNA libraries were generated using the Illumina DNA Prep kit following the manufacturer’s instructions. Libraries were then pooled and sequenced on an Illumina NovaSeq 6000 platform with 2 x 150 bp. On average, samples yielded 8.8 ± 2.7 (mean ± SD) million reads. Both 16S amplicon sequencing and shot-gun sequencing were performed at the Competence Centre for Genomic Analysis (Kiel).

### 16S rRNA amplicon sequencing data preprocessing

MiSeq 16S amplicon sequence data were processed using DADA2 workflow (Callahan *et al*., 2016) (https://benjjneb.github.io/dada2/bigdata.html) resulting in abundance tables of amplicon sequence variants (ASVs). Taxonomy was assigned using the Bayesian classifier provided in DADA2 and using the Silva rRNA database v.138 (Quast *et al*., 2013). Samples with > 5000 reads were retained for analyses. After this step, the 16S rRNA gene amplicon sequencing of the main and validation experiments retained on average 20K ± 8K and 10K ± 4K (mean ± SD) reads per sample, respectively. ASVs not assigned to Bacteria or annotated as Mitochondria, Chloroplast, or Eukaryote were removed. DADA2 outputs were combined into a single object and phyloseq R (1.40.0) (McMurdie and Holmes, 2013) package was used for downstream analyses.

### Shot-gun metagenomics data preprocessing

Shot-gun reads were processed using an in-house established workflow from our institute (https://github.com/ikmb/TOFU-MAaPO) that relies on BBtools and bowtie2 for mapping and host decontamination, and bioBakery tools (MetaPhlAn 4.0 and HUMaN 3.6) for taxonomic and functional potential profiling. Raw reads were quality trimmed and mapped to the mouse genome (GRCm39) to discern between sequenced reads from the host and the gut microbiome bacteria. Only samples containing > 1 Gbp after trimming and host decontamination were kept in the analysis.

Taxonomic features and quantification of microbial communities’ relative abundances on the 72 samples were done by using MetaPhlAn 4.0 (Manghi *et al*., 2023) with default parameters and the custom SGB database. MetaPhlAn 4.0 relies on unique marker genes of 26,970 species-level genome bins (SGBs) from a collection of highly curated 1.01M prokaryotic and metagenome-assembled genomes (MAGs). Likewise, functional potential profiling (stratified pathways, gene families, and enzyme categories) in the same samples was computed using HUMAnN 3.6 (Beghini *et al*., 2021), with default parameters. All the abundances were transformed to CPMs before any statistical analysis.

### 16S and metagenomics downstream analyses

Most of the univariate and multivariate analyses for the 16S and metagenomics data were done in the R statistical software (v.4.2.1) under phyloseq (McMurdie and Holmes, 2013) (v.1.40.0), vegan (Dixon, 2003) (v.2.6-2), variancePartition (Hoffman and Schadt, 2016)(v.1.26.0) and MAasLin2 (Mallick *et al*., 2021) (v.1.10.0).

Within sample diversity (alpha diversity) was explored by computing diversity indexes (Chao1, Shannon, Simpson, Observed ASVs) on ASV abundance data and finding differences among certain groups (Wilcoxon rank-sum test or Kruskal-Wallis tests were performed).

Between-sample diversity (beta diversity) and the differences between maternal genotype groups and time were explored and visualized in a Principal Coordinate Analysis (PCoA) plot. It was quantified as Aitchison distances on centered-log ratio transformed ASVs counts.

Associations of microbiome composition to maternal genotype or other covariates were tested with the implementation of PERMANOVA models (using the adonis2 function from the vegan package). The P and R^2^ values were determined by 10,000 permutations using time. R^2^ was used for computing partial omega squares (Olejnik and Algina, 2003; Lakens, 2013) as an unbiased estimator of effect sizes. P values were subject to Benjamini-Hochberg correction.

### Temporal shifts and genotype-specific differential abundance analysis

To detect differences in changes of microbial features (16S rRNA, metagenomics) and pathways (metagenomics), between the maternal genotype over time, we built linear mixed models on center-log transformed abundances in the MaAslin2 package (Mallick *et al*., 2021) that included time, maternal genotype, and individual mice as a random variable (*microbial abundances* ∼ (1|*m ice*) + *maternal genotype* + *time*). P values were corrected for multiple hypothesis testing using the Benjamin-Hochberg procedure, and a false discovery rate < 0.05 was defined as the significant threshold. Features (taxa, or pathways) appearing in at least 80% of the samples per group were included in the analysis. For the analysis of temporal dynamics, we built linear mixed-models for each maternal genotype (*Atg16l1^fl/fl^*and *Atg16l1^ΔIEC^*) data: *microbial abundances* ∼ (1|*mice*) + *time* to identify genera that significantly changed the abundance at any time point (week 3 or week 6) compared with BL.

### Variance Partition Analysis

To understand the major sources of variation in our fecal microbiota data at different phylogenetic resolutions (Phylum, Order, Class, Family, Genus and ASVs), functional categories (Gene Ontology, KEGG Orthology, Enzyme Comission, MetaCyc Reactions), and metabolic pathways (MetaCyc Pathways), we applied variancePartition (v.1.26.0) (Hoffman and Schadt, 2016), which uses linear mixed models to compute the attributable percentage of variation of a feature based on selected covariates. Here, we selected Individual mouse, maternal genotype, time, lipocalin, maternal weight, and number of pups as possible contributors to the variation of the fecal microbiota composition.

### Data and Code Availability

The 16S rRNA amplicon sequencing and the shot-gun metagenomics data are accessible through the European Nucleotide Archive (ENA, https://www.ebi.ac.uk/ena accessed on 8th November 2023) under the accession number PRJEB70999. Additional data supporting the findings of this study, as well as all codes used to generate the bioinformatic analyses, are available from the corresponding author and will be made available through Github.

## Results

### Longitudinal differences of fecal microbiota of *Atg16l1^fl/fl^* and *Atg16l1^ΔIEC^* during pregnancy and lactation periods

To explore the functional impact of *ATG16L1-*mediated autophagy in the intestinal epithelium on the dynamics of fecal microbiota during pregnancy, we performed 16S rDNA and shotgun sequencing in 15 weeks female littermate villin (V)-cre+; *Atg16l1^fl/fl^* (hereafter called *Atg16l1^ΔIEC^*) or villin (V)-cre^-^ *Atg16l1^fl/fl^* (hereafter called *Atg16l1^fl/fl^* or WT ) mice (Aden *et al*., 2018) . We used sequential sample from the day before timed mating (baseline, termed BL), from trimester 3 (at the end of week 3 after timed mating, termed w3) and from the end of lactation (at the end of week 6, termed w6) (Fig. 1A). , In the 16S rRNA data, significant differences in α−diversity (within-sample) between *Atg16l1^fl/fl^*and *Atg16l1^ΔIEC^* animals were neither observed during pregnancy nor at the end of lactation (Fig. 1B, Supp. Fig 1A). In contrast, microbiome community composition (β-diversity, between-sample), varied significantly between *Atg16l1^fl/fl^* and *Atg16l1^ΔIEC^* in the third trimester (w3) compared to BL (representing the third trimester PERMANOVA on between-sample Aitchison distances, adj.R2 = 0.063, Benjamini-Hochberg corrected p = 2.9e^-4^) and recovered until the end of the lactation period (w6) (Fig. 1C, Supp. Fig 1B, 1C, and 1D).

**Figure 1.**
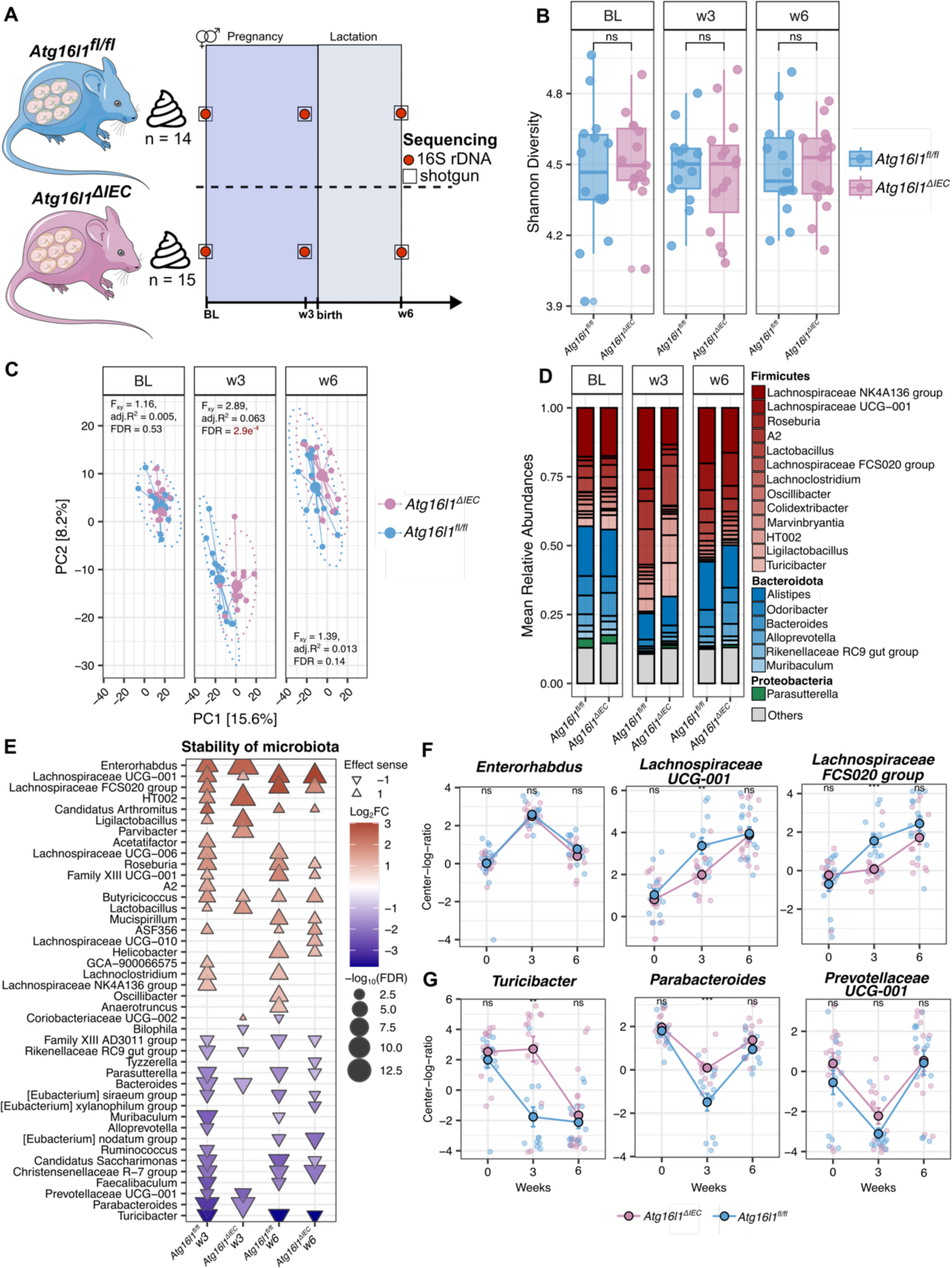
Gut microbiota characterization of *Atg16l1^fl/fl^* and *Atg16l1^ΔIEC^*mice during pregnancy and after lactation periods. **A.** Study design. Stool samples of *Atg16l1^fl/fl^* (n = 14) and *Atg16l1^ΔIEC^*(n = 15) pregnant mice were collected at baseline (before mating), week 3 (late pregnancy), and week 6 (after lactation period). Samples were submitted to 16S rRNA and shotgun sequencing. **B.** Alpha diversity analysis of gut microbiota during pregnancy and lactation. Shannon diversity was computed at the ASV level. Cross-sectional comparisons between *Atg16l1^fl/fl^*and *Atg16l1^ΔIEC^* were performed using the Wilcoxon rank-sum test. **C.** Principal coordinate analysis on Aitchison distance matrix of pregnant mice. Differences between *Atg16l1^fl/fl^* and *Atg16l1^ΔIEC^* were tested with PERMANOVA with 10000 permutations. FDR represents Benjamini-Hochberg corrected *p* values, and adj.R^2^ represents partial omega squares as effect size in the analysis of variance. **D.** Relative abundances of the top 20 most abundant genera. Unclassified genera and those with low relative abundance are grouped as “Others”. Colors represent individual Phylum and color gradients represent individual genus within a Phylum. **E.** Triangular dot-plot showing significantly changing genera compared to baseline. The triangles show the hues and direction of the effect size (log_2_FC). Color intensity and size represent the magnitude of the effect size and FDR significance of each specific genus, respectively. **F.** Top three unstable genera with significantly increased abundances in any timepoint compared to baseline. **G.** Top three unstable genera with significantly decreased abundances in any timepoint compared to baseline. The longitudinal plots are colored to remark differences in genotype.

At the taxonomic level of most abundant taxa, we found – as expected-(Koren *et al*., 2012) an overall pregnancy-related increase in mean relative abundances of Firmicutes and a reduction of Bacteroidetes that seem to recuperate after the lactation period in both *Atg16l1^fl/fl^*and *Atg16l1^ΔIEC^* mice (Fig 1D). Interestingly, appreciable differences were observed between *Atg16l1^fl/fl^* and *Atg16l1^ΔIEC^* mice at w3 for some genera, e.g., a significant decrease in *Lachnospiracea NK4A136 group* and *Lachnospiraceae UCG-001* and a significant increase in *Turicibacter* in *Atg16l1^ΔIEC^* mice (Fig 1D).

To further understand the dynamics and stability of certain taxa during pregnancy and lactation, we performed a multivariate analysis using MaAsLin2 (Mallick *et al*., 2021) using the 16S rRNA data. We built linear mixed models to identify which genera showed a time-dependent significant difference in abundance compared to BL in *Atg16l1^fl/fl^* and *Atg16l1^ΔIEC^* mice. We found 42 temporal shifts of genera that increased or decreased significantly in abundance at w3 (trimester 3) and/or w6 (after lactation) compared to BL (FDR < 0.05, Fig 1E, Supp. Fig 2A and 2B).

At the end of pregnancy (w3), 6 genera, belonging to the Actinobacteriota (*Enterorhabdus*) and Firmicutes (*Lachnospiraceae* UCG-001, *Lactobacillaceae* HT002, *Ligilactobacillus*, *Butyricicoccus* and *Lactobacillus*) phyla, significantly increased in both, *Atg16l1^fl/fl^* and *Atg16l1^ΔIEC^*, whereas 4 genera, belonging to Bacteroidota (*Rikenellaceae* RC9 gut group, *Bacteroides*, *Prevotellaceae* UCG-001, and *Parabacteroides*) phyla, significantly decreased in both, *Atg16l1^fl/fl^* and *Atg16l1^ΔIEC^*.

The same comparison at the end of the lactation period (w6), showed that 9 genera, mostly belonging to Firmicutes (*Lachnospiraceae* UCG-001, *Lachnospiraceae* FCS020 group, *Candidatus Arthromitus*, *Roseburia*, *Family XIII* UCG-001, *Butyricicoccus*, *Clostridium sp. ASF356*), Campylobacteria (*Helicobacter*), and Deferribacteres (*Mucispirillum*) phyla increased in both, *Atg16l1^fl/fl^*and *Atg16l1^ΔIEC^*, whereas 7 genera, belonging to Firmicutes (*Family XIII AD3011* group, *[Eubacterium] siraeum* group, *[Eubacterium] nodatum* group, *Christensenellaceae* R-7 group and *Turicibacter*), Proteobacteria (*Parasutterella*), and Patescibacteria (*Candidatus Saccharimonas*) phyla, significantly decreased in both, *Atg16l1^fl/fl^* and *Atg16l1^ΔIEC^*. Although the abundance shift direction (up or down) was similar for numerous genera in both groups, we noted a diminished count of genera demonstrating a significant temporal shift in *Atg16l1^ΔIEC^*mice compared to *Atg16l1^fl/fl^* mice and several genera showed a significant difference between the groups at the timepoints w3 and/or w6 (Fig. 1E).The top three most unstable genera by effect size (Log_2_FC) in *Atg16l1^fl/fl^* and *Atg16l1^ΔIEC^* animals are represented in Fig 1F and Fig 1G. We could confirm several genera with common longitudinal dynamics by mining shotgun metagenomics data at strain-level resolution (Supp. Fig 2C and 2D). Specifically, at w3, *Lactobacillus johnsonii*, *Limosilactobacillus reuteri,* and *Ligilactobacillus murinus* significantly increased and *Muribaculum gordoncarteri* and *Muribaculaceae bacterium* Isolate 110 HZI decreased in abundances (Supp. Fig 2E). In summary, the results suggest that loss of functional *Atg16l1* in the intestinal epithelium of mice is associated with a perturbation of pregnancy-induced fecal bacterial community shifts during pregnancy.

### Maternal genotype is the main driver of explainable fecal microbiota variation at the end of pregnancy between *Atg16l1^fl/fl^* and *Atg16l1^ΔIEC^* mice

We next studied the relative contribution of an ATG16L1 deletion in comparison to other experimental factors to overall microbiota variation. To methodically investigate the impact of experimental covariates, we conducted a variance partition analysis (Hoffman and Schadt, 2016) to quantify the extent to which the variation in each taxon can be attributed to differences in specific covariates. This analysis was done using the 16s rRNA data on different phylogenetic levels (Phylum, Class, Order, Family, Genus, and ASVs) and four different comparisons: (1) a complete dataset including samples from all time points to account for time-dependent effects, and on subsets of samples representing the three individual time points, i.e. (2) only baseline, (3) only w3 and (4) only w6. Before building the variance partition models, we identified co-linearity between the different experimental covariates by canonical correlation analysis (Supp. Fig 3A). Those covariates with a correlation coefficient < 0.6 were considered non-colinear and therefore were included in the variance partition models. Variance partition models were built for the respective data sets in the following way; (1) all time points : *Taxa Abundances* ∼ *maternal genotype + weeks + number of pups + lipocalin* ; and (2) each time point subset: *Taxa Abundances ∼ maternal genotype + maternal weight + number of pups + lipocalin* (Supp. Fig 3).

The complete model could explain around 31% of the total variation (Fig 2A and Supp Table S2). Specifically, the main driver of variation of the fecal microbiota across samples was time (approximately 26% in all taxa), followed by the maternal genotype (approx. 2.4%), number of pups (approx. 1.92%), and lipocalin (around 1.0%) (Supp Table S2). Also for the time point subsets, BL and w6 (at the end of lactation), the maternal genotype had a small (approx. 2%) contribution to the microbial variation; interestingly, the maternal genotype was the main driver of variation in the trimester 3 timepoint (w3) with around 14%, followed by the number of pups (approx. 6 %), lipocalin ( around 4 %) and maternal weight (around 3%) (Supp Table S2), in line with the previous observation for the β−diversity analysis.

**Figure 2.**
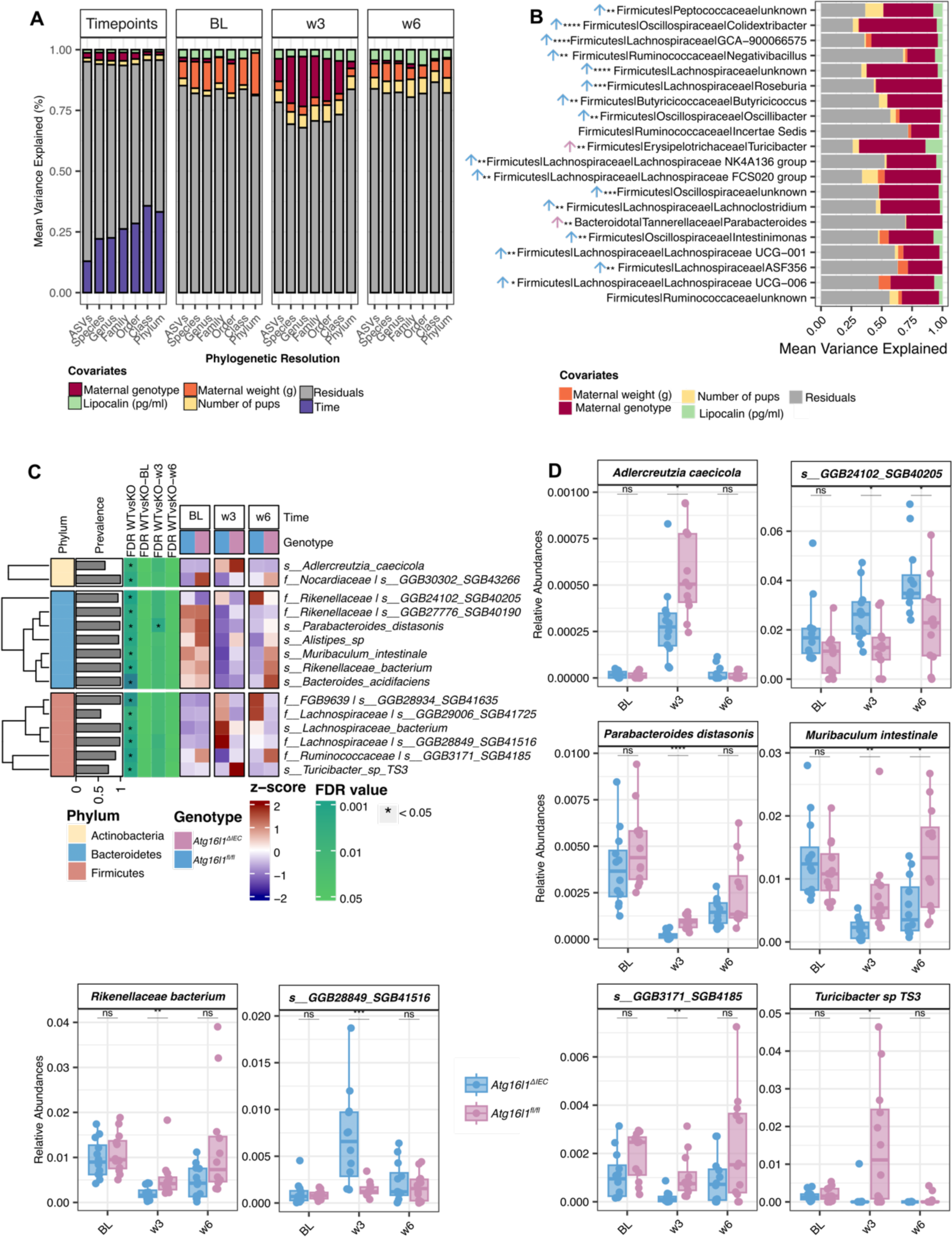
Taxonomic profiling differences of the gut microbiota of *Atg16l1^fl/fl^* and *Atg16l1^ΔIEC^* during pregnancy and lactation periods. **A.** Variation partition analysis of different taxa-level abundances from 16S rDNA data. Each color represents the contribution of individual covariates as a source of variation. Each stacked area bar plot compartment (Timepoints, BL, w3, and w6) represents the output of different variance partition models. **B.** Top 20 genera mainly contributing to the variation of the maternal genotype at week 3. Cross-sectional comparisons (*Atg16l1^fl/fl^* vs *Atg16l1^ΔIEC^*) were performed using the Wilcoxon rank-sum test and p values were corrected by the Benjamini-Hochberg method. *: *p* < 0.05 , **: *p* < 0.01, ***: *p* < 0.001, ****: *p* < 0.0001, empty: not significant. The color of the arrow symbol represents the sense of the increased effect of each significant genus (*Atg16l1^fl/fl^*, blue or *Atg16l1^ΔIEC^*, pink). **C.** Heatmap of significant abundance changes in identified species and SGBs found during multivariate comparison of *Atg16l1^fl/fl^* vs *Atg16l1^ΔIEC^*. For each cell, colors indicate the row-wise z-score of relative abundances, asterisks denote the FDR < 0.05 significance at each cross-sectional comparison and prevalence represents the percentage of non-zero features. Row-wise clusters represent features that belong to the same Phylum. **D**. Box plots showing the relative abundance of selected species and SGBs.

We then examined the top 20 genera that contribute the most to the variation associated with maternal genotype in trimester 3 (Fig. 2B). Notably, among the top 20 ranked features, we found a broad spectrum of genera, including Firmicutes , such as *Colidextribacter* (VE = 64%), *Turicibacter* (VE = 55%), *Roseburia* (VE = 54%), *Lachnoclostridium* (VE = 49%), various types of *Lachnospiraceae* (VE = 46%, VE = 40%, VE = 36%, and VE = 30%), and Bacteroidota like the genus *Parabacteroides* (VE = 30%), which together explain most of the genotype-dependent differential abundances between pregnant *Atg16l1^fl/fl^* and *Atg16l1^ΔIEC^* mice in trimester 3 (week 3) (full list in Supplementary Tables S3 and S4).

### Species-level differences in the fecal microbiota of pregnant and lactating *Atg16l1^fl/fl^* and *Atg16l1^ΔIEC^* mice

We next investigated species-level (and ASV) differences between the fecal microbiota of *Atg16l1^fl/fl^* and *Atg16l1^ΔIEC^*mice using shotgun metagenomics (and 16S rRNA) data. We again performed a multivariate analysis to extract genotype-dependent significant differences in species abundance at the specified time points (BL, w3, and w6). We built four types of linear mixed models in MaAsLin2; a complete model: *bacteria abundances ∼ (1|individual mice) + time + maternal genotype* and three models for each time point (BL, w3 and w6): *bacteria abundances ∼ maternal genotype*, accounting for the maternal genotype and in the complete model for the effect of time and individual trajectories of mice.

We found 15 metagenomic features that were significantly different between *Atg16l1^fl/fl^ and Atg16l1^ΔIEC^*. From these, only 8 were identified at species level, and 7 were identified as species-level genome bins (SGBs). Specifically, we found 6 Firmicutes (*Turicibacter sp. TS3*, and 4 SGBs belonging to *Lachnospiraceae* and *Ruminoccaceae* families), 7 Bacteroidetes (*Parabacteroides distanosis*, *Alistipes sp*., *Muribaculum intestinale*, *Bacteroides acidifaciens*, and 4 SGBs belonging to *Rikenellaceae* family), and 2 Actinobacteria (*Adlercreutzia caecicola* and 1 SGB belonging to *Nocardiaceae* family). Like in the genus level analysis, most of the species and SGBs had similar longitudinal overall abundance patterns (up or down) between genotypes, but many of the taxa differed significantly in relative abundance at the end of pregnancy (w3) (Figure 2C and 2D). For instance, *Adlercreutzia caecicola, Muribaculum intestinale, Parabacteroides distanosis, Turicibacter sp. TS3,* and *Rikenellaceae* bacterium exhibited a notable rise in relative abundance in *Atg16l1^ΔIEC^* compared to *Atg16l1^fl/fl^* at w3. In contrast, certain SGBs associated with the *Rikenellaceae* family (SGB40205) and *Lachnospiraceae* family (SGB41516) displayed the opposite pattern. We could confirm that similar taxa profiles were present at ASV level (Supp. Fig 4A and B).

### Inference of functional differences in the fecal microbiota of *Atg16l1^ΔIEC^* and *Atg16l1^fl/fl^* mice from shotgun metagenomics

Having demonstrated that genetic deletion of ATG16L1 in the intestinal epithelium in mice is associated with altered trajectories of fecal microbiota changes during and after pregnancy, we next aimed to understand the underlying functional consequences using metagenomic pathway profiling.

We inferred the presence and abundance of microbial community functions using the HUMAnN 3.6 pipeline (Beghini *et al*., 2021) from the metagenomics data. A total of 494799 gene families and 324 metabolic pathways were detected. Gene families were regrouped into 4 functional categories: KEGG Orthogroups (KEOrs), Level-4 enzyme commission (EC) categories, Gene Ontology (GO), and MetaCyc Reactions (RXN). Similarly, metabolic pathways were classified into 4 different superclasses and subclasses according to MetaCyc database (Karp *et al*., 2019; Caspi *et al*., 2020). The contribution of different experimental covariates to the functional potential of the fecal microbiota was again analyzed by variance partition analysis (Hoffman and Schadt, 2016) with the same data subsets as in the taxonomical analyses.

The complete model explained around 30 % of the total variation (Fig 3A, detailed values for each individual category in Supp. Table S6), with high concordance across the 4 functional categories (KEOr, EC, GO, RXN). Similar to our observations at the taxonomic level, the primary factor influencing the variation in fecal microbiota was time (approx. 26% in all categories), succeeded by the number of pups (around 1.3%), maternal genotype (around 1%), and lipocalin (around 0.7%) (Supp. Table S6). The percentage of variance explained by the maternal genotype to around 4% at week 3 compared to BL (approx. 1.7%), week 6 (approx. 2 %) (Fig. 3A and Supp Table S6). For all functional levels tested, the increase, however, was less pronounced compared to the taxonomical analysis.

**Figure 3.**
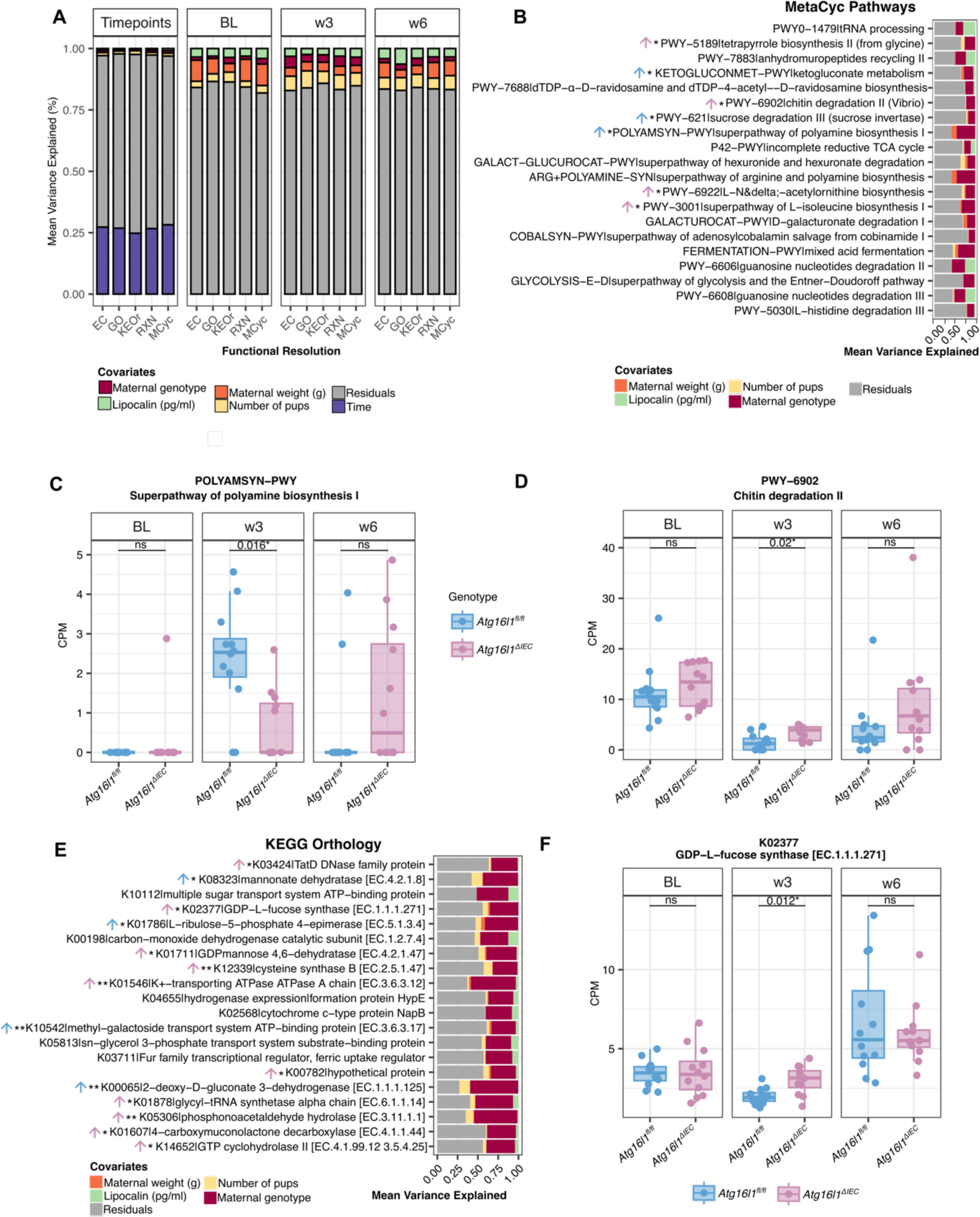
Functional potential predictions of the gut microbiota of *Atg16l1^fl/fl^* and *Atg16l1^ΔIEC^* during pregnancy and lactation periods. **A.** Variation partitioning analysis of different functional categories abundances from metagenomics data. Each color represents the contribution of individual covariates as a source of variation. Each stacked-area bar plot compartment (Timepoints, BL, w3, and w6) represents the output of different variance partition models. **B.** Top 20 MetaCyc pathways mainly contributing to the variation of the maternal genotype at week 3. **C.** Boxplot of the abundances of Polyamine biosynthesis pathway. The color of the arrow symbol represents the sense of the increased effect of each significant genus (*Atg16l1^fl/fl^*, blue or *Atg16l1^ΔIEC^*, pink). **D.** Boxplot of the abundances of Chitin degradation pathway. Boxplots are colored to remark differences in genotype. **E.** Top 20 KEGG orthologs that most contribute to the variation of maternal genotype at week 3. The arrow symbol and colors represent the sense of the increased effect of each significant genus (*Atg16l1^fl/fl^*, blue or *Atg16l1^ΔIEC^*, pink). **F.** Box plot of the K02377 gene family, GDP-L fucose synthase. Boxplots are colored to remark differences in genotype. Cross-sectional comparisons (*Atg16l1^fl/fl^* vs *Atg16l1^ΔIEC^*) were performed using the Wilcoxon rank-sum test and p values were corrected by the Benjamini-Hochberg method. *: *p* < 0.05 , **: *p* < 0.01, ***: *p* < 0.001, ****: *p* < 0.0001, empty: not significant.

Interestingly, we found strong variation (variance explained > 20%) in the MetaCyc pathways associated with maternal genotype at week 3 (Fig 3B and Supp. Table S7). The alterations were positioned in key metabolic pathways e.g. Amine and Polyamine biosynthesis (POLYAMSYN-PWY: superpathway of polyamine biosynthesis I [VE = 45%, FDR < 0.05] and ARG+POLYAMINE-SYN: superpathway of arginine and polyamine biosynthesis [VE = 45%, FDR < 0.05]). (Supp Table S7, Supp Table S8).

A formal test for differential abundance (FDR < 0.05) between *Atg16l1^fl/fl^*and *Atg16l1^ΔIEC^* animals for each pathway, (see methods for details), verified seven of the top 20 pathways. Notably, polyamine biosynthesis functions (ARG+POLYAMINE and POLYAMSYN-PWY) emerged as the most diminished function in *Atg16l1^ΔIEC^* mice at week 3, with the lowest FDR recorded at 0.016. (Fig 3C, Supp. Table S8). Furthermore, we also observed a significant reduction of the term biosynthesis of the nucleotide sugar dTDP-α-D-ravidosamine (PWY-7688), ketogluconate metabolism (KETOGLUCONMET-PWY), and sucrose degradation III (PWY-621) in *Atg16l1^ΔIEC^*mice compared to *Atg16l1^fl/fl^*, whereas we observed significantly increased abundances on amino acid biosynthesis (L-isoleucine and acetylornithine), chitin degradation (PWY-6902) and tetrapyrrole biosynthesis (PWY-5189). On the KEGG level (Fig 3D and Supp. Table S9) terms (KEOrs) associated with genotype-dependent variation at week 3 included e.g., Pentose and glucuronate interconversions (K00065: *kduD*; 2-deoxy-D-gluconate 3-dehydrogenase [VE=59%, FDR < 0.01], and K08323: *rspA*; mannonate dehydratase [VE=43%, FDR < 0.05]), and KEOrs associated to Amino sugar and nucleotide sugar metabolism (K01711: *gmd*; GDPmannose 4,6-dehydratase [VE=37%, FDR < 0.05], and K02377: *fcl*; GDP-L-fucose synthase [VE = 35%, FDR < 0.05]). Again, a formal test for differential abundance of the top 20 KEOrs, confirmed 14 to be significantly different between *Atg16l1^fl/fl^*and *Atg16l1^ΔIEC^* mice at timepoint w3 (FDR < 0.05) (Fig 3E, F and Supp. Table S10). This included KEOrs from the Phosphonate and phosphinate metabolism (K05306: *phnX*; phosphonoacetaldehyde hydrolase) and the two-component system (K01546: *kdpA*; K^+^-transporting ATPase ATPase A chain) and the Amino sugar and nucleotide sugar metabolism (K01711: *gmd*; GDP mannose 4,6-dehydratase, and K02377: *fcl*; GDP-L-fucose synthase), which were confirmed as significantly increased gene families in *Atg16l1^ΔIEC^*mice at week 3. Most significantly decreased abundances on KEOrs were associated to Pentose and glucuronate interconversions (K00065: *kduD*; 2-deoxy-D-gluconate 3-dehydrogenase, and K08323: *rspA*; mannonate dehydratase), followed by a KEOr of ABC transporter (K10542: *mglA*; methyl-galactoside transport system ATP-binding protein) , and Pentose phosphate pathway (K01786: L-ribulose-5-phosphate 4-epimerase).

### A validation experiment recapitulates fecal microbiota differences between *Atg16l1^ΔIEC^* and *Atg16l1^fl/fl^* in trimester 3

The previous fecal microbiota characterization suggested that the main effect of the genetic ablation of *ATG16L1* in intestinal epithelial cells can be observed at the end of pregnancy (week 3). To confirm this finding, we performed a validation experiment consisting of a weekly collection and sequencing (16S rDNA) of stool samples throughout timed pregnancy, from BL to w3 (Fig. 4A). At the end of pregnancy, mice were sacrificed, and their biological material was used for phenotypic analysis, including pro-inflammatory markers and characterization of pubs. Results were compared to the respective pregnant littermate wildtype *Atg16l1^fl/fl^*.

**Figure 4.**
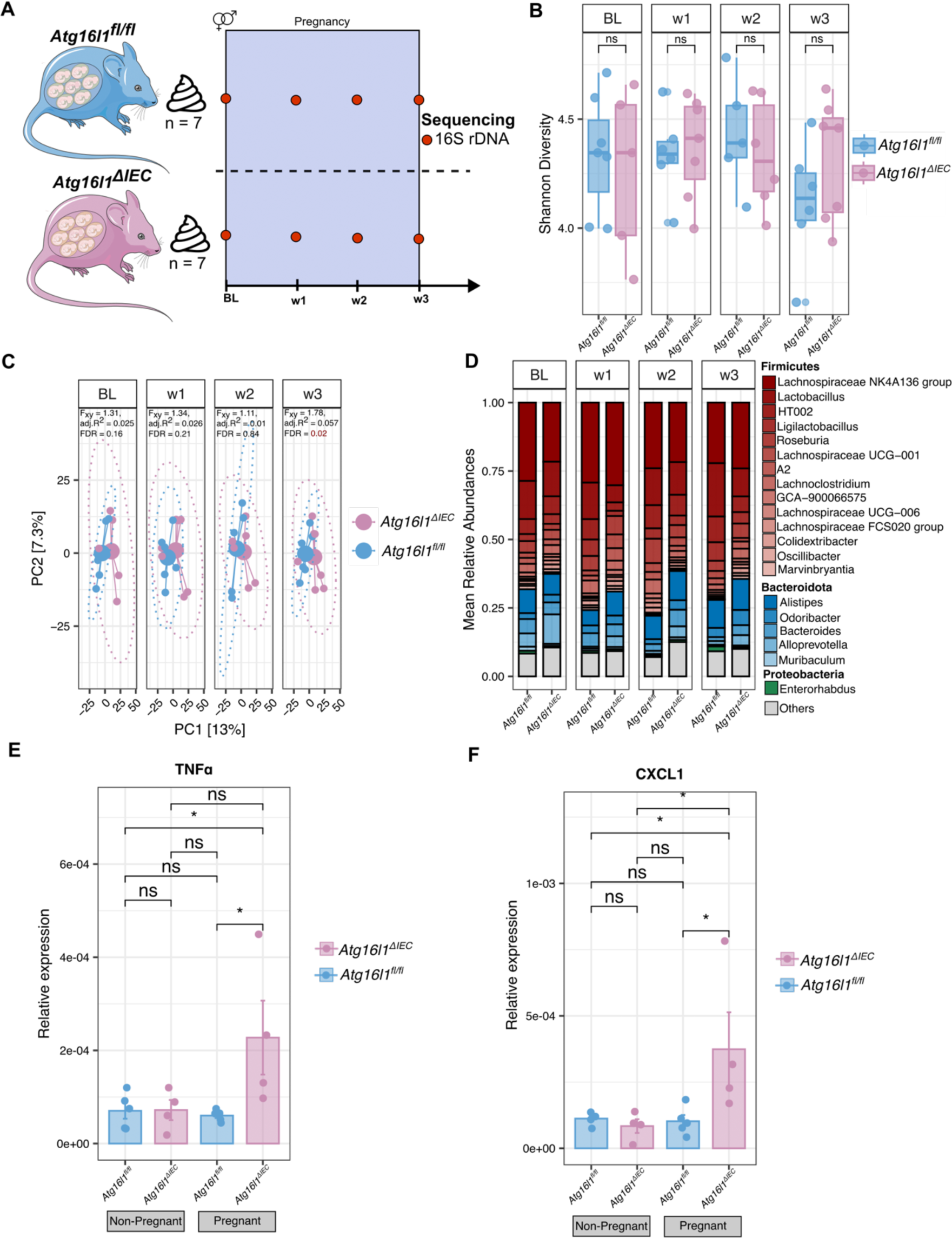
Gut microbiota and cytokine expression characterization of *Atg16l1^fl/fl^* and *Atg16l1^ΔIEC^* during pregnancy period (independent validation experiment). **A.** Study design. Stool samples of *Atg16l1^fl/fl^*(n = 7) and *Atg16l1^ΔIEC^*(n = 7) pregnant mice were collected weekly from BL (before mating) to week 3 (late pregnancy) and submitted to 16S rRNA amplicon sequencing. **B.** Alpha diversity analysis of gut microbiota during pregnancy and after lactation. Shannon diversity was computed at the ASV level. **C.** Principal coordinate analysis on Aitchison distance matrix of pregnant mice. Differences between *Atg16l1^fl/fl^*and *Atg16l1^ΔIEC^* were tested with PERMANOVA with 10000 permutations. FDR represents Benjamini-Hochberg corrected *p* values, and adj.R^2^ represents partial omega squares as effect size in the analysis of variance. D. Relative abundances of the top 20 most abundant genera. Unclassified genera and those with low relative abundance are grouped as “Others”. Colors represent individual Phylum and color gradients represent individual genus within a Phylum. **E, F.** Expression levels of the cytokines in nulliparous and pregnant *Atg16l1^fl/fl^* and *Atg16l1^ΔIEC^* mice. **E.** *TNFα* and **F.** *CXCL1* expression by Taqman assay. Groups are shown as nulliparous *Atg16l1^fl/fl^*, nulliparous *Atg16l1^ΔIEC^,* Pregnant *Atg16l1^fl/fl^* and Pregnant *Atg16l1^ΔIEC^*. Cross-sectional comparisons (*Atg16l1^fl/fl^* vs *Atg16l1^ΔIEC^*) were performed using the Wilcoxon rank-sum test. *: *p* < 0.05, **: *p* < 0.01, ***: *p* < 0.001, ****: *p* < 0.0001, ns: not significant.

During pregnancy period, no significant differences or trend of changes in within-sample diversity were observed between *Atg16l1^fl/fl^* and *Atg16l1^ΔIEC^*(Fig. 4B). In addition, between sample diversity varied between *Atg16l1^fl/fl^*and *Atg16l1^ΔIEC^* only at w3 (PERMANOVA on between-sample Aitchison distances, adj.R2 = 0.057, Benjamini-Hochberg corrected p = 0.02) (Fig. 4C). These results are aligned to the microbial composition we observed in the previous experiment. (Fig 1B, and 1C).

Exploring the top 20 most abundant genera, we again found an overall pregnancy-related increase of mean relative abundances of Firmicutes and an overall reduction of Bacteroidetes in both, *Atg16l1^fl/fl^* and *Atg16l1^ΔIEC^* mice (Fig 4D). Although no formally significant differences were observed between *Atg16l1^fl/fl^* and *Atg16l1^ΔIEC^* mice at week 3 at the genus level, we observed a clear trend towards a decrease in *Lachnospiracea NK4A136 group*, *Lachnospiraceae UCG-001*, *Lachnoclostridium*, *Lactobacillus* and *Roseburia* in *Atg16l1^ΔIEC^* mice (Fig 4D, Supp. Fig 5A) that recapitulates the findings in Fig 1D.

Interestingly, the differential abundance analysis at the ASV level showed significant differences in ASVs from the Firmicutes and Bacteroidetes phylum between *Atg16l1^fl/fl^* and *Atg16l1^ΔIEC^* during pregnancy (Supp. Fig 5B). Specifically, we saw significant differences at week 3 in two *Lachnospiraceae* family members as *Lachnospiraceae NK4A136 group*, and *Eubacterium xylanophilum group*; and, in the *Eubacterium brachy group*, an *Anaerovoracaceae* family member.

### *Atg16l1* deletion in IECs results in increased inflammatory cytokine secretion in the colonic mucosa of pregnant mice in trimester 3

To understand the physiological and phenotypic effects caused by the absence of *Atg16l1* on intestinal epithelial cells and its consequences for pregnancy, we compared different molecular and physiological parameters between nulliparous *Atg16l1^ΔIEC^* (BL) animals and pregnant *Atg16l1^ΔIEC^* (week 3) animals (Supp. Fig 6). As expected, pregnant (week 3) mice showed increased body weight compared to nulliparous (BL) mice within the same genotype: *Atg16l1^ΔIEC^* (*p* < 0.05) and *Atg16l1^fl/fl^* (*p* < 0.01) (Supp. Fig 6A), while there was no detectable difference between the groups at timepoint w3.

Other parameters (liver, spleen weight and colon length) presented without significant changes between genotypes as well as timepoints tested (Supp. Fig 6B, 6C and 6E). Of note, caecum weight was significantly higher in pregnant *Atg16l1^ΔIEC^*mice compared to pregnant *Atg16l1^fl/fl^* (*p <* 0.05) (Supp. Fig 6D). Interestingly, we found that pups born from *Atg16l1^ΔIEC^* mice were significantly lighter than those that were born from *Atg16l1^fl/fl^*mothers (*p* < 0.01) (Supp. Fig 6F). However, no significant differences were found in their length (Supp. Fig 6G). To further explore the potential impact of changes in microbiota composition during pregnancy on the host, we measured levels of several cytokines in the colonic mucosa of pregnant mice by qPCR. We discovered significantly elevated mRNA levels of pro-inflammatory cytokines, including *Tnf-α* (p < 0.05) and *Cxcl1* (p < 0.05), in pregnant *Atg16l1^ΔIEC^* mice compared to *Atg16l1^fl/fl^* mice (Fig 4E and 4F), suggesting that the deletion of *Atg16l1* in intestinal epithelial cells leads to increased pregnancy-induced intestinal inflammation, potentially originating from the rise of pro-inflammatory bacteria due to the depletion of autophagy and antimicrobial functions in the intestine.

## Discussion

Pregnancy is associated with a plethora of transient alterations in maternal organ systems, which are all geared toward accommodating the requirements of fetal growth and development. These alterations also comprise structural and compositional changes in the microbiota at different body sites (Edwards *et al*., 2017; Amabebe and Anumba, 2021), which are thought to provide an important contribution to the overall adaptation for maximizing reproductive success, e.g. by providing additional energy from nutrient intake (Koren *et al*., 2012). Relatively little is known about how the host shapes this physiological remodelling of the resident microbial communities, particularly in the gut, and how these changes could contribute to the observed immune alterations during pregnancy.

Intestinal Paneth cells are pivotal players in mucosal immunity and stem cell maintenance and contribute to maintaining a homeostatic intestinal microbiome. We hypothesized that antimicrobial effector functions of the intestinal epithelium could be among the factors that shape the observed pregnancy-associated changes in the intestinal microbiota. Our data from genetically modified mice now provides evidence for a role of the Crohńs disease risk gene *ATG16L1* in intestinal epithelial cells (IECs) in this process. Hypomorphic or genetically deleted *ATG16l1* in the IEC lineage in mice (and humans) has been shown to cause impaired antimicrobial Paneth and goblet cell architecture (Cadwell *et al*., 2008), to increase the propensity to necroptotic cell death (Matsuzawa-Ishimoto *et al*., 2017) and to impair responses to the protective cytokine Il-22 (Aden *et al*., 2018) , which is known to regulate anti-microbial effectors, e.g. defensins in IECs. We have previously demonstrated that under steady state conditions the microbiota of *Atg16l1^ΔIEC^* mice and their respective floxed littermate controls are nevertheless surprisingly similar and that changes in the microbiota only arise from additional perturbations, e.g. when challenging the mice with DSS (Tschurtschenthaler *et al*., 2017). We thus chose this model as an entry point to understand the effect of pregnancy as a physiological perturbation of the intestinal microbiota.

Our mouse model experiment enabled us to study the dynamics of gut microbiota transitioning between non-pregnancy and pregnancy states (week 0 to trimester 3), and then returning to a non-pregnant state (week 3 to week 6). In control and *Atg16l1^ΔIEC^* mice, we observed deep changes in β-(between-sample) diversity with time, more specifically, at the timepoint week 3, which corresponds to the third trimester or pre-partum. This finding is consistent with other studies showing an increased distance in β-diversity of fecal communities during pregnancy, culminating shortly before delivery (Koren *et al*., 2012). α−(within sample) diversity indices of fecal microbiota stayed constant during gestational time in our experiments adding to the controversy on these measures during pregnancy in rodent models and humans. While Koren et al. showed a decrease of within-sample species richness, which contributed to coining of term gestational dybiosis (Koren *et al*., 2012), other publications did not find temporal changes of α-diversity during pregnancy (Faas *et al*., 2020). In line with our results, a recent large human study involving approximately 1,500 pregnant Chinese women found no notable variation in fecal α−diversity throughout the entire pregnancy period (Yang *et al*., 2020). We observed a significant rise in Firmicutes and a decrease in the Bacteroidetes phyla, which confirmed earlier findings in pregnant mice (Faas *et al*., 2020). Human studies have reported more variable changes already on the phylum level, from a slight increase of *Firmicutes* phylum (Liu *et al*., 2017; Selma-Royo *et al*., 2021) to dominant changes in *Actinobacteria*, and *Proteobacteria* (Koren *et al*., 2012). The heterogeneity of maternal fecal microbiota changes in humans has been attributed to recommended or habitual (e.g. increase of carbohydrates) diet changes (Mandal *et al*., 2016; Röytiö *et al*., 2017; Gomez-Arango *et al*., 2018; Kunasegaran *et al*., 2023).

What were the main alterations of fecal bacterial communities during pregnancy in the absence of functional *Atg16l1* in the intestinal epitheliumStarting from a similar baseline composition, we observed a significant β-diversity difference between *Atg16l1^fl/fl^* and *Atg16l1^Δ^*^IEC^ littermate female mice at the end of trimester 3 (week 3), which almost returned to normal at the end of lactation (week 6). Interestingly, we show that control (i.e. *Atg16l1^fl/fl^*) mice had more systematic temporal changes in individual genera compared to *Atg16l1^ΔIEC^*mice. Thus, it can be speculated that normal epithelial function including the secretion of antimicrobial effector molecules such as defensins is involved in orchestrating the dynamics of different taxa at the end of pregnancy. This is similar to the situation in female reproductive tract, where regulation of defensins modulates the succession of the resident microbiota during gestation.

The main taxonomical differences between *Atg16l1^fl/fl^* and *Atg16l1^ΔIEC^* mice comprised members of the *Lachnospiraceae*, *Bacteroidaceae*, *Ruminnococcaceae,* and *Muribaculaceae* families. Specifically, we found a marked reduction of the Short-Chain Fatty Acid (SCFA) producing *Lachnospiraceae* family members like *Roseburia*, *Lachnospiraceae UCG-001*, *Lachnospiraceae NK4A136*, *Lachnoclostridium*, *Lachnospiraceae UCG-006*, and *Lachnospiraceae FCS020 group* and an increase in bacteria like *Turicibacter sp*, *Parabacteroides distasonis*, *Bacteroides acidifaciens*, *Muribaculum instestinale*, *Adlercreutzia caecicola* and *Ruminococcus sp* in *Atg16l1^ΔIEC^*mice during pregnancy. The same taxonomical abundance pattern was found for *Lachnospiraceae*, *Bacteroides ovatus*, and *Parabacteroides* in male mice carrying the *Atg16l1* T300A polymorphism before and after Dextran Sulfate Sodium-induced inflammation, and stool transplantation of active CD patients into germ-free mice (Lavoie *et al*., 2019).

Interestingly, many taxa that we found as increased or decreased in *Atg16l1^ΔIEC^* pregnant mice in trimester 3 have been previously described to have a role in intestinal inflammation, specifically in Crohn’s disease. For instance, a significant reduction in the SCFA-producing bacteria *Roseburia* has been a consistent pattern of CD (Takahashi *et al*., 1960; Shen *et al*., 2018, 2022; Ma *et al*., 2022), while the role of other members of the *Lachnospiraceae* family is more controversial (Vacca *et al*., 2020). Of note, *Lachnospiraceae UCG-001*, the same genus that we found reduced in *Atg16l1^ΔIEC^*, has been negatively associated with CD in a recent study (Liu *et al*., 2022). *Bacteroides acidifaciens* has been characterized as a symbiotic organism that inhibits obesity, enhances insulin sensitivity in mice (Yang *et al*., 2017), which however, was also associated to gut inflammation in mice models of colitis (Berry *et al*., 2015; Staley *et al*., 2023). *Parabacteroides*, which had a higher abundance in *Atg16l1^ΔIEC^* pregnant mice at week 3 compared to controls, were found to be highly abundant in early CD patients (Ma *et al*., 2022) and specifically *Parabacteroides distasonis* has been identified in intestinal microlesions of a severe CD patient (Yang *et al*., 2019). Also, this bacterium was reported to induce depressive behavior in a mouse model of CD-like ileitis (Gomez-Nguyen *et al*., 2021).

The role of *Turicibacter* and *Adlercreutzia* on gut inflammation is less clear. *Turicibacter sanguinis* has been isolated from a febrile patient with acute appendicitis (Bosshard, Zbinden and Altwegg, 2002). In a mouse model with depletion of CD8+ T cells, *Turicibacter* was found increased in abundance (Presley *et al*., 2010), while it was found low in mice lacking TNF expression prior colitis induction (Jones-Hall, Kozik and Nakatsu, 2015). In contrast to our results, Adlercreutzia spp. have been found at lower abundance in CD patients and in relatives of patients with CD (Shaw *et al*., 2016; Leibovitzh *et al*., 2022). These discrepancies could be associated with the fact that *Adlercreutzia caecicola*, the species we found increased in *Atg16l1^ΔIEC^* mice at the end of pregnancy, was originally isolated from the cecal content of male C57BL/6 mice (Stoll *et al*., 2021). Further studies on *Turicibacter* and *Adlercreutzia* genus are necessary to elucidate its role in IBD and gut inflammation.

In addition, inference of metabolic properties from shot-gun metagenomics data revealed that the gut microbiota alterations of *Atg16l1^ΔIEC^*mice at the end of pregnancy also influence their metabolic capacity. Particularly, we identified reduced capacity to biosynthesize polyamines, metabolize acidic polysaccharides like keto-gluconate and degradate sucrose. Although little is known about the role of microbial-derived polyamines like putrescine and spermidine, recent studies have shown that dysbiosis in gut microbiota can trigger imbalances in polyamine metabolism which may favour metabolic-related diseases (Tofalo, Cocchi and Suzzi, 2019). In fact, spermidine has been shown to exert a protective role against experimental colitis in mice. Supplementation with spermidine suppressed colitis and reduced colitis-associated carcinogenesis (Gobert *et al*., 2022). On the other hand, we identified that *Atg16l1^ΔIEC^*mice have an improved capacity to produce tetrapyrroles (like vitamin B12), branched-chain amino-acid L-isoleucine, and has a higher capacity to produce fucosylated oligosaccharides (via GDP-L-fucose synthase), this last function has been found in fucose-carrier *Bacteroides* species of Crohn’s disease patients (Kappler *et al*., 2020). These changes were associated with a spontaneous inflammatory response in the colonic mucosa of *Atg16l1^ΔIEC^*mice at trimester 3 as evidenced by elevated mRNA levels of *Tnfa* and *Cxcl1* and decreased weight of offspring compared to pregnant control mice. Importantly, the observed subtle inflammatory response occurred in the colon and well outside of the time window of the previously reported spontaneous inflammation in the mouse strain which had an onset around the age of 35 weeks (Tschurtschenthaler *et al*., 2017).

Limitations of the study comprise a relatively small size of the validation cohort, which does recapitulate all features of the exploratory cohort. In addition, we do not provide formal proof, e.g. by using germ-free *Atg16l1^ΔIEC^*mice, that the observed physiological changes between *Atg16l1^ΔIEC^*mice and their respective *Atg16l1^fl/fl^* littermate controls are indeed causally related to the microbial changes. Hence, the study warrants a deeper investigation using a gnotobiotic setup to understand the role of the identified taxa and their associated metabolites for a healthy pregnancy outcome. Our observation regarding the functional potential of the fecal microbiota relies on metagenomic data, it will thus be important to investigate the inferred changes on the metabolomic level to derive potential targets for metabolic biomarkers and/or therapeutic intervention ( e.g. by supplementing spermidine as a protective postbiotic).

In summary, our study suggests that genetic impairment of autophagy in IECs may amplify the pro-inflammatory tone at late stages of gestation (Koren *et al*., 2012). *Atg16l1^ΔIEC^* pregnant mice have overall lesser coordinated changes in their fecal microbiota than their wildtype littermate controls. Several of the salient changes present in *Atg16l1^ΔIEC^*female mice in trimester 3 resemble features observed in human IBD patients, e.g. reduced propensity for spermidine production. It is thus tempting to speculate that these changes in female IBD patients with genetic variants in autophagy-related genes such as ATG16L1 T300A (Hampe *et al*., 2007; Rioux *et al*., 2007; Lavoie *et al*., 2019) might modulate the threshold of inflammatory relapses around birth.

Our results highlight the importance of better understanding host factors contributing to the functional shifts of the intestinal ecosystem in IBD during gestation and after birth. Clearly, it underscores the necessity for larger-scale cohort studies in female IBD patients to capture the individual heterogeneity and distinct sources of variation of host-microbe interactions in pregnancy. We suggest that rationale microbiota-targeted therapies may represent a powerful addition to improve personalized care and pregnancy outcomes in female patients with IBD.

## Ethics approval

All animal experiments were approved by the local animal safety review board of the federal ministry of Schleswig Holstein and conducted according to national and international laws and policies (V 241 - 69128/2016 (3-1/17) and V 242-7224.121-33)

## Consent for publication

Not applicable.

## Availability of data and materials

All data generated or analyzed during this study are included in this published article and its supplementary information files.

## Competing Interests

The authors declare that the research was conducted in the absence of any commercial or financial relationships that could be construed as a potential conflict of interest.

## Funding

This work was supported by the Deutsche Forschungsgemeinschaft (DFG) Research Unit 5042 “miTarget” (TP 03, Grant number RO 2994/10-1), Collaborative Research Center (CRC) 1182 “Origin and Function of Metaorganisms” (subprojects C02 and Z03, Grant number CRC 1182/1 and 2), Cluster of Excellence 306 “Inflammation at Interfaces” (Grant number EXC 306/2) and the Bundesministerium für Bildung und Forschung (BMBF) funded e:Med grant project “iTREAT: A systems medicine approach to Individualized TREATment trajectories in Psoriasis and IBD” (subproject SP5, Grant number 01ZX1902A)

## Authors’ contributions

V.A.L-A, M.F., A.K. and P.R, conceived and designed experiments. M.F., and A.L. performed the animal experiments, and V.A.L-A, M.F., A.R. and R.B. analyzed the animal and sequencing data. F.S., A.R., R.B., E.W., and D.E. supported the preprocessing of the sequencing data. V.A.L-A., M.F., and P.R. wrote the manuscript, and all authors revised the manuscript for important intellectual content and approved the final version. P.R. supervised this work.

## Acknowledgments

The authors would like to thank Vivian Wegener (IKMB, Kiel) for the support with the sample amplification and library preparation of the 16S data, and Dr. Christina Bronowski (Liverpool University) for the discussion and feedback about the 16S rRNA and metagenomics analyses. We also thank the Competence Centre for Genomic Analysis (CCGA), Kiel, Germany, where the 16S rRNA amplicon sequences and shot-gun metagenomics were performed.

## Supplementary Figure Legends

**Supplementary Figure 1.**
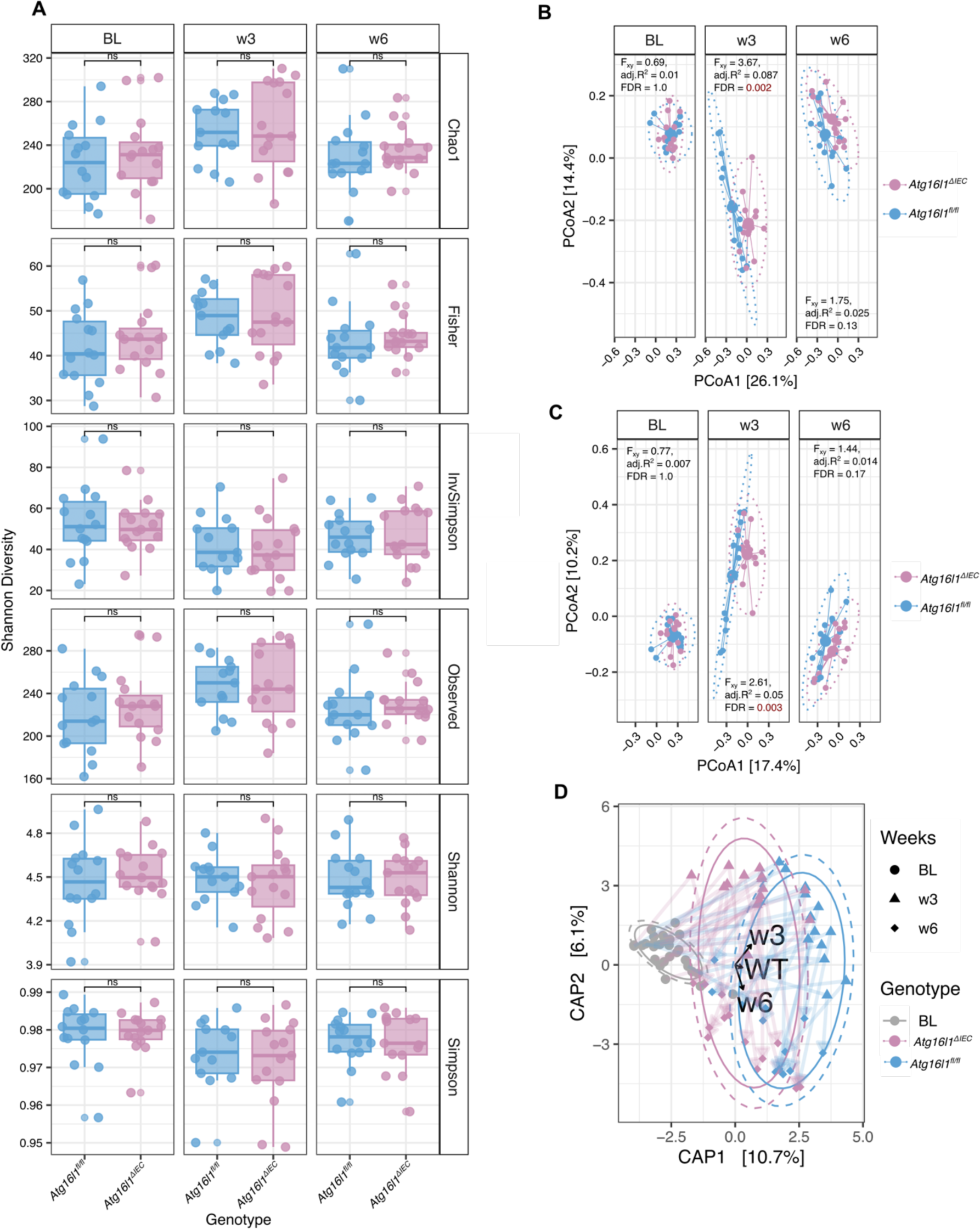
Microbial composition differences between *Atg16l1^fl/fl^* and *Atg16l1^ΔIEC^*. **A.** Within-sample diversity indexes. **B.** Principal Coordinate Analysis from Bray-Curti’s distance matrix. **C.** Principal coordinate analysis from Jaccard distance matrix. **D.** distance-based redundancy analysis on Aitchison distances. Time and Genotype were used as association variables. Differences in alpha diversity between *Atg16l1^fl/fl^* and *Atg16l1^ΔIEC^* were performed using the Wilcoxon rank-sum test, *: *p* < 0.05, **: *p* < 0.01, ***: *p* < 0.001, ****: *p* < 0.0001, ns: not significant. Differences in beta diversity between *Atg16l1^fl/fl^* and *Atg16l1^ΔIEC^*were tested with PERMANOVA with 10000 permutations. FDR represents Benjamini-Hochberg corrected *p* values, and adj.R^2^ represents partial omega squares as effect size in the analysis of variance

**Supplementary Figure 2.**
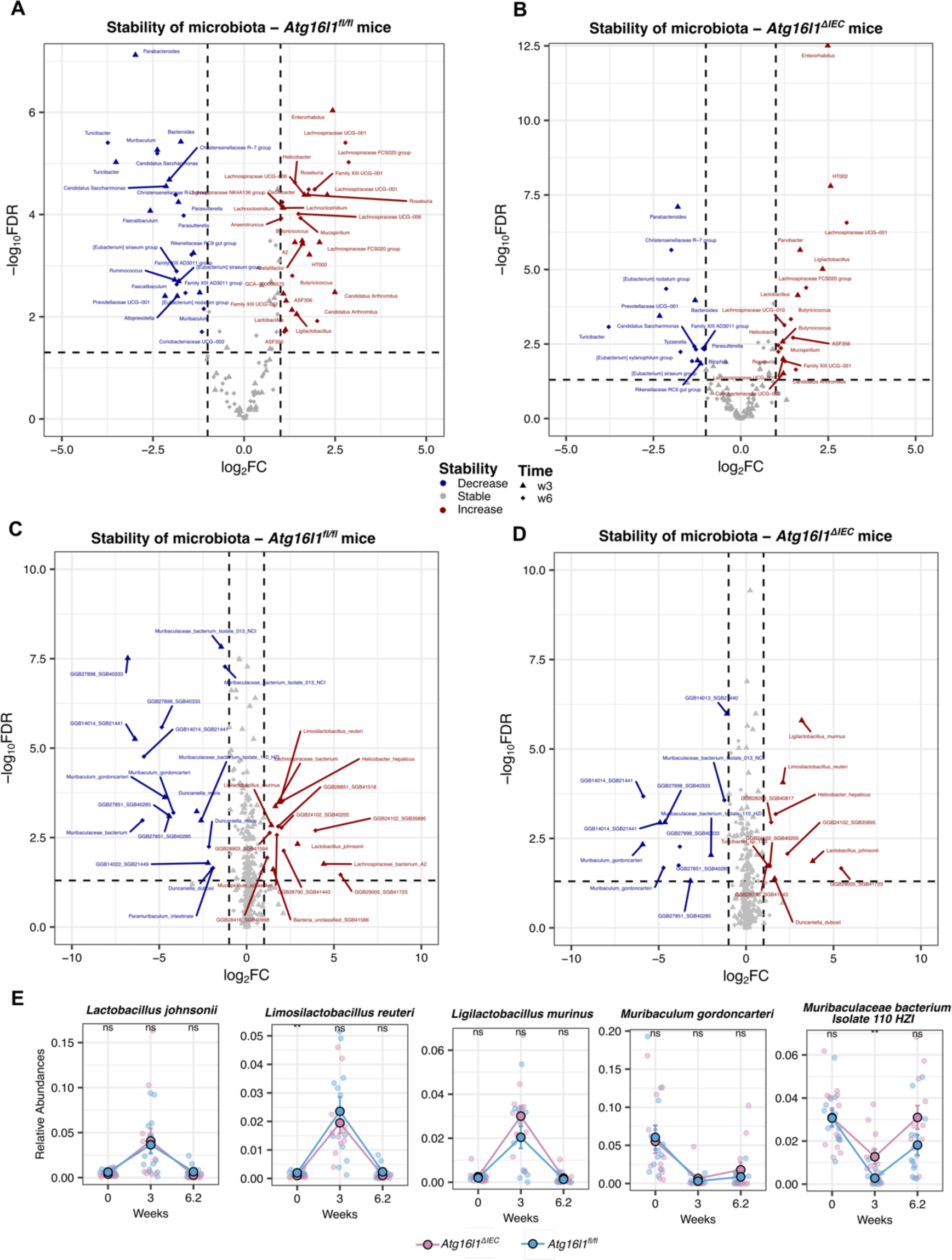
Temporal dynamics analysis. **A.** Volcano plot depicting the most changing genera (|Log2FC| > 1 and FDR < 0.05) in *Atg16l1^fl/fl^* pregnant mice from 16S rDNA data. Genus in red or blue represent those genera that increase or decrease in abundance in at least one time point, respectively. **B.** Volcano plot depicting the most changing genera (|Log2FC| > 1 and FDR < 0.05) in *Atg16l1^ΔIEC^* pregnant mice from 16S rDNA data. **C.** Volcano plot depicting the most changing species (|Log2FC| > 1 and FDR < 0.05) in *Atg16l1^fl/fl^* pregnant mice from shot-gun metagenomics. **D.** Volcano plot depicting the most changing species (|Log2FC| > 1 and FDR < 0.05) in *Atg16l1^ΔIEC^* pregnant mice from shot-gun metagenomics. E. Species with similar temporal dynamics in both *Atg16l1^fl/fl^* and *Atg16l1^ΔIEC^*.

**Supplementary Figure 3.**
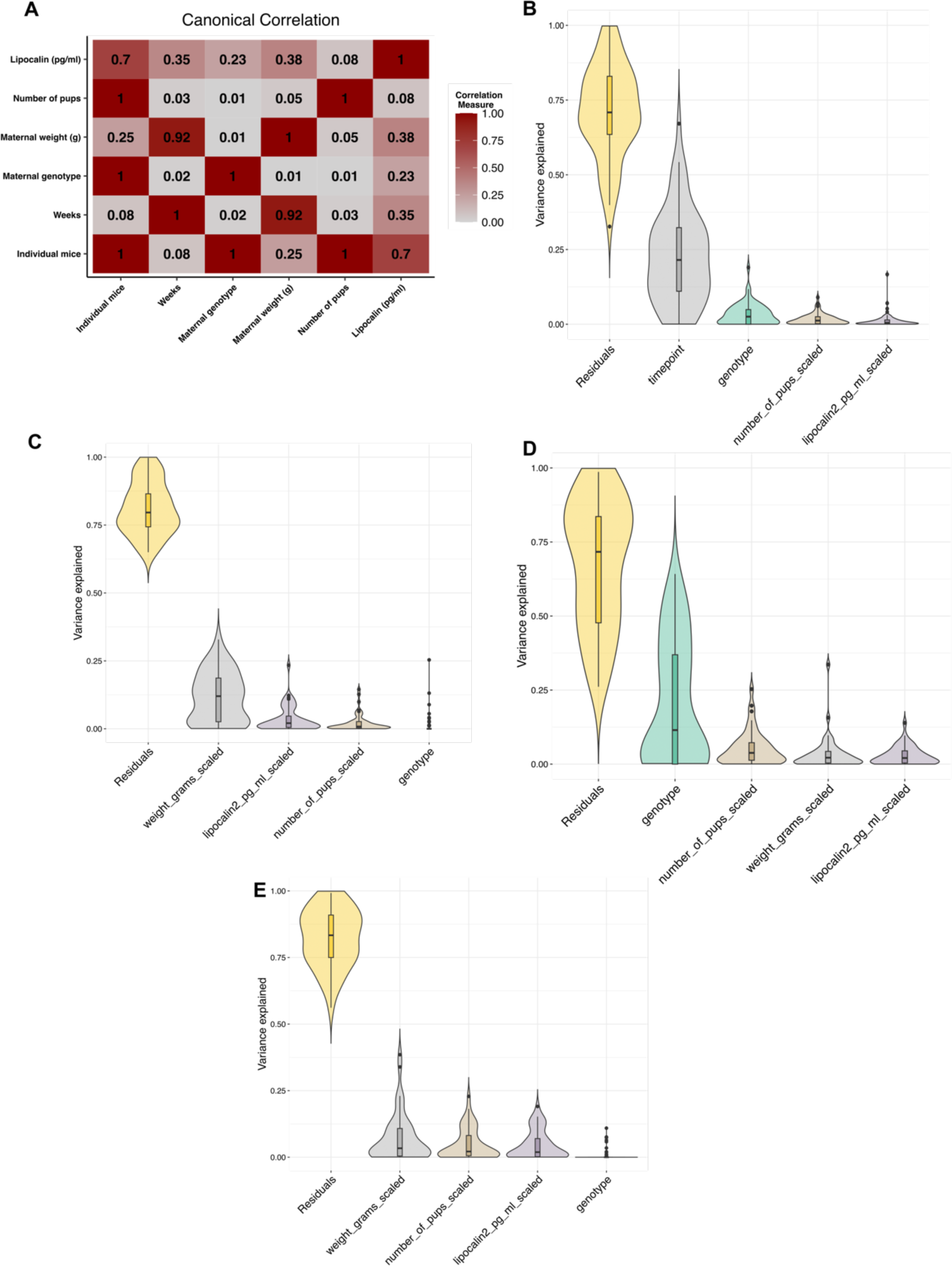
Variance partition analysis on 16S rDNA amplicon sequencing data. **A.** Canonical correlation analysis between covariates used in variancePartition models. Covariates-pairs with correlation coefficient > 0.6 are considered co-linear and were deleted from the models. **B.** Variation contribution plot of the complete dataset variance partition model for genus features. **C.** Variation contribution plot of the baseline model for genus features. **D.** Variation contribution plot of the week 3 variance partition model for genus features. E. Variation contribution plot of the week 6 variance partition model for genus features.

**Supplementary Figure 4.**
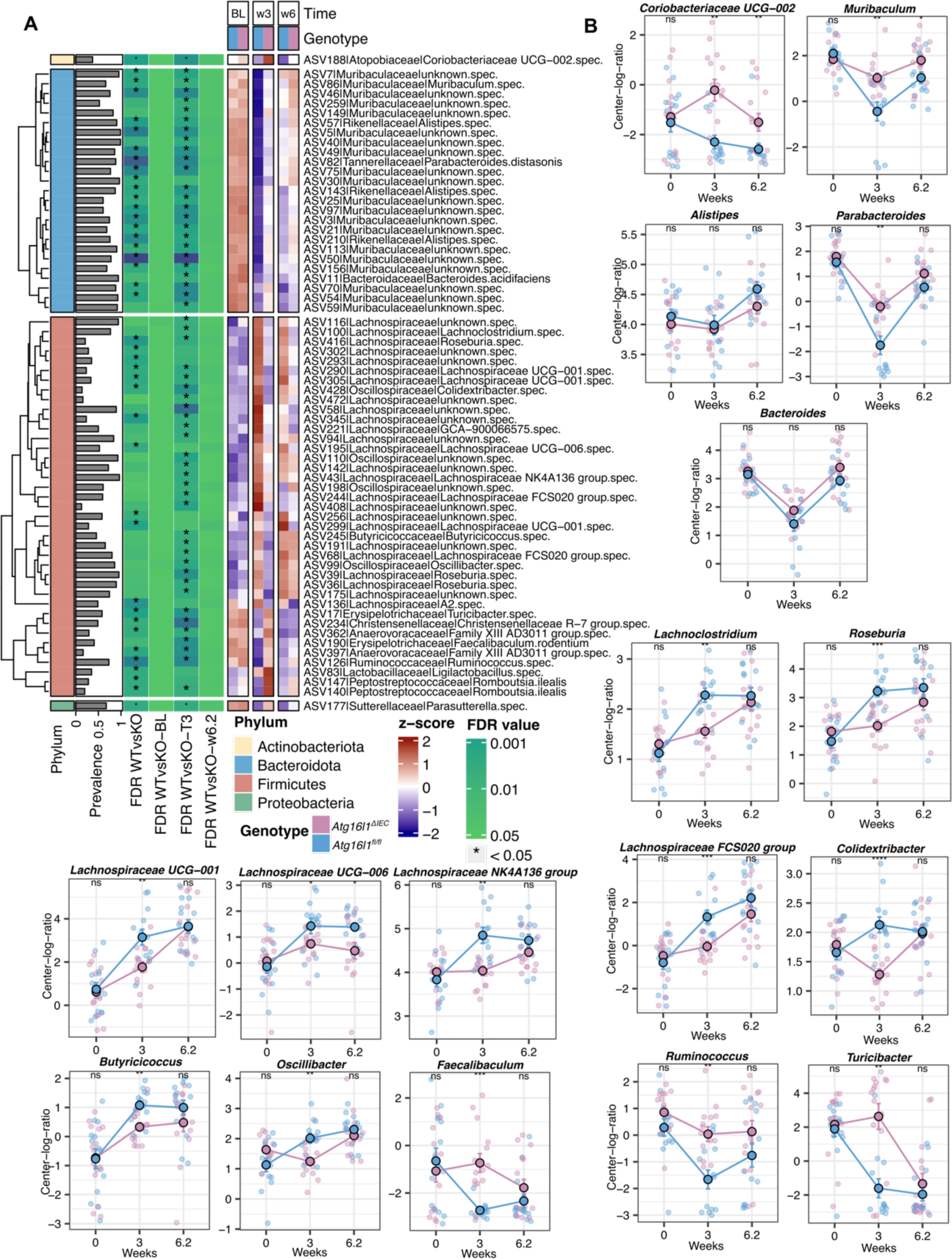
Differential abundance analysis at ASV level. **A.** Heatmap of significant abundance changes in ASVs found during the linear mixed model comparison between *Atg16l1^fl/fl^* vs *Atg16l1^ΔIEC^*. For each cell, colors indicate the row-wise z-score of the center-log-ratio transformed mean abundances, asterisks denote the FDR significance at each cross-sectional comparison and prevalence represents the percentage of non-zero features. Row-wise clusters represent features that belong to the same Phylum. **B.** longitudinal plots of different genera that were identified at Supp. Figure 4A.

**Supplementary Figure 5.**
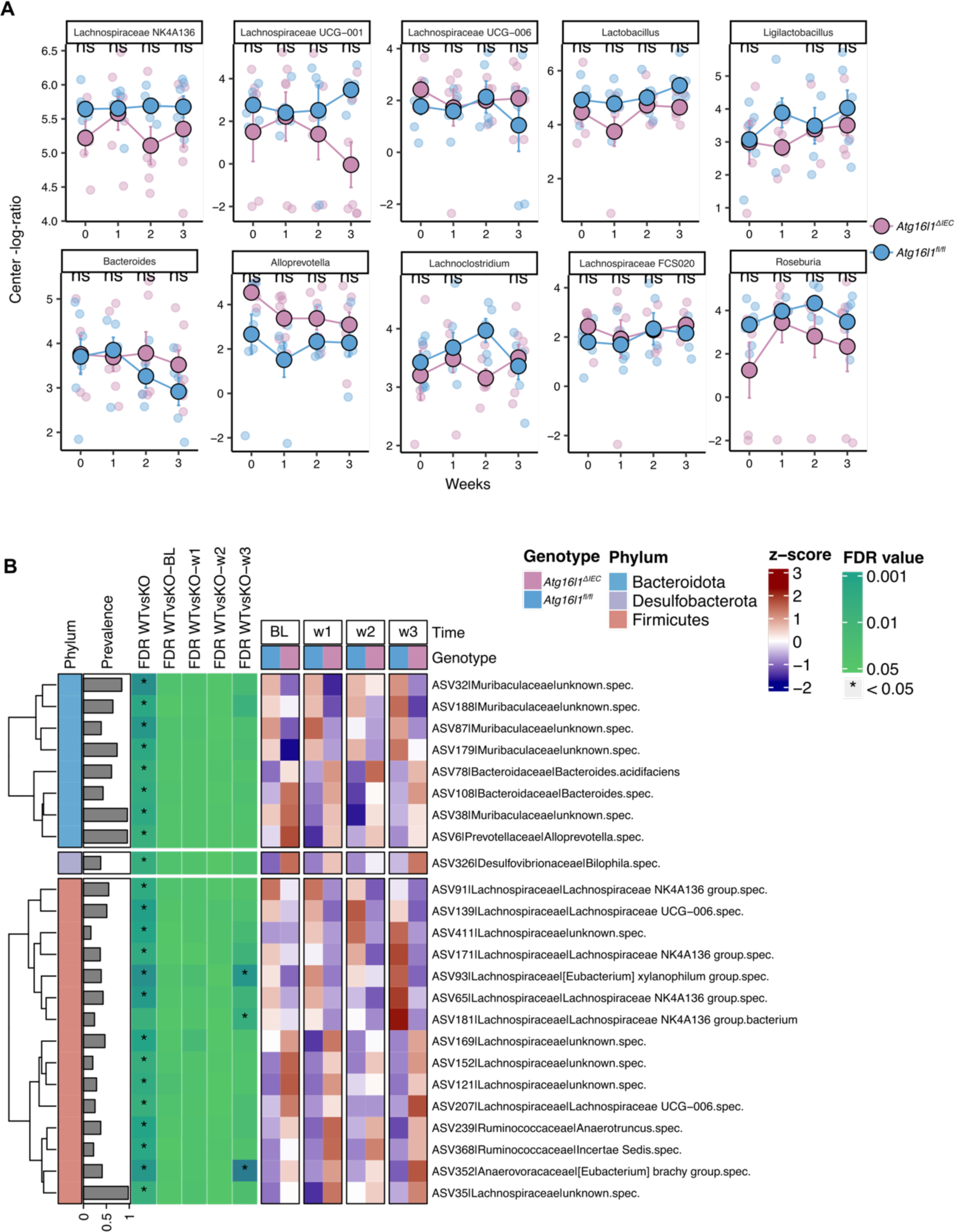
**A.** Longitudinal center-log ratio transformed abundances of selected taxa during pregnancy (validation experiment, 16S rDNA data). **B.** Heatmap of significant abundance changes in ASVs found during the linear mixed model comparison between *Atg16l1^fl/fl^* vs *Atg16l1^ΔIEC^* pregnant mice from the validation experiment. For each cell, colors indicate the row-wise z-score of center-log ratio transformed mean abundances, asterisks denote the FDR significance at each cross-sectional comparison and prevalence represents the percentage of non-zero features. Row-wise clusters represent features that belong to the same Phylum.

**Supplementary Figure 6.**
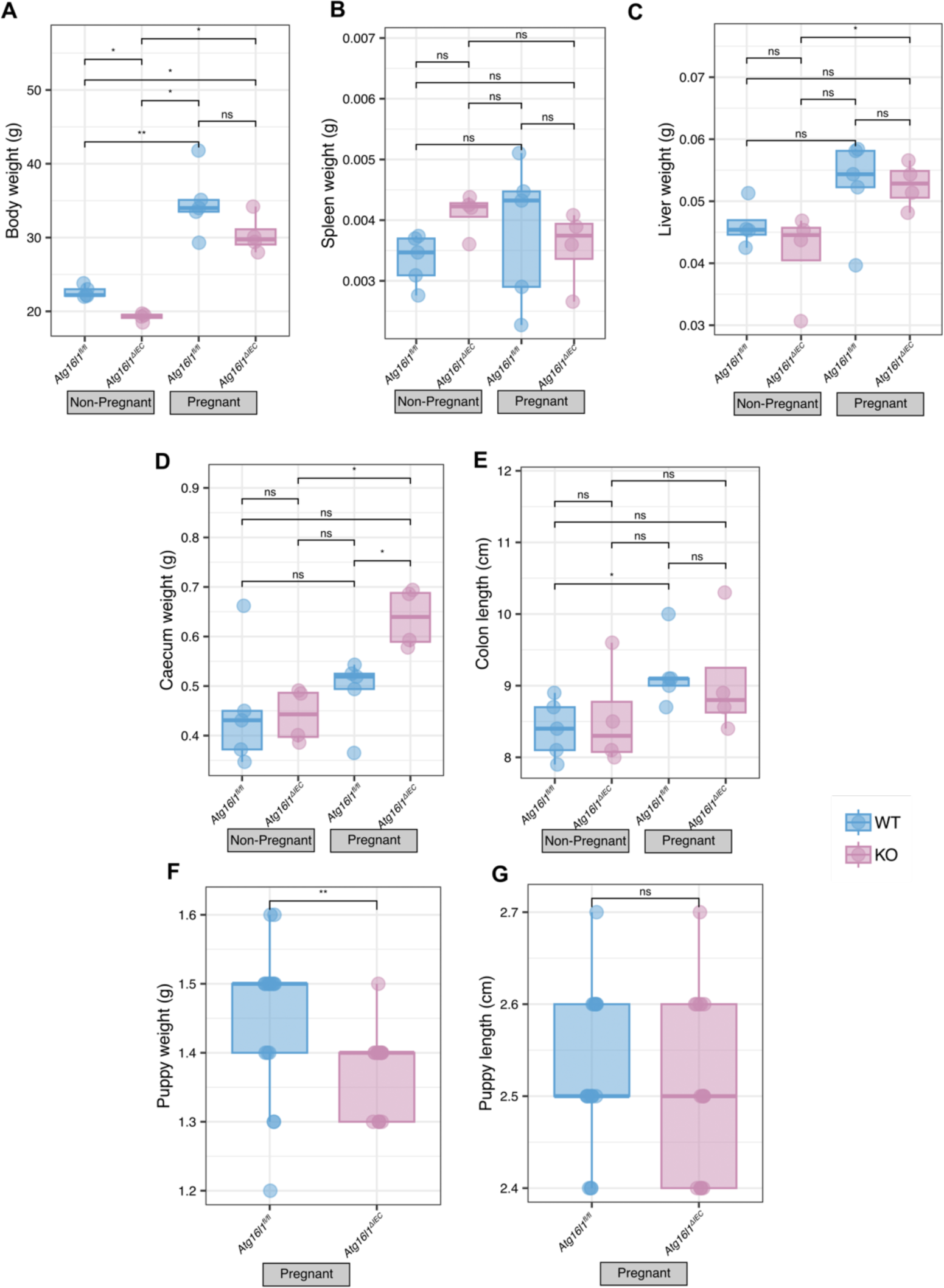
Histological phenotypes for nulliparous *Atg16l1^fl/fl^*, nulliparous *Atg16l1^ΔIEC^*, pregnant *Atg16l1^fl/fl^*and pregnant *Atg16l1^ΔIEC^* showing. **A.** Body weight. **B.** Spleen weight. **C.** Liver weight **D.** Caecum weight **E.** Colon length. **F.** Puppy weight. **G.** Puppy length. The significance of differences was determined using the nonparametric Mann Whitney U test * *P*<0.05.

## Notes

### Competing Interest Statement

The authors have declared no competing interest.

